# Unlocking the Genetic Potential of *Solanum bulbocastanum* (SB22, Selection 22): A Valuable Resource for Enhancing Disease Resistance in Commercial Potato Cultivars

**DOI:** 10.1101/2024.03.20.586016

**Authors:** Senthilkumar Shanmugavel, Kelly Vinning, Sam C. Talbot, Charles R. Brown, Vidyasagar Sathuvalli

## Abstract

Cultivated potatoes are susceptible to a host of diseases caused by various pathogens. Wild relatives of potatoes are used in breeding programs as sources of resistance introgressed into cultivated potatoes. The wild potato *Solanum bulbocastanum* is an essential source of resistance to Columbia root knot nematode (CRKN) and late blight. We present the initial chromosome-level assembly of SB22, produced using PacBio long reads and Dovetail Hi-C scaffolding. The final assembly size was 655.3 Mb. Using the BRAKER pipeline, 43,280 gene models were predicted, with a BUSCO completeness of about 90.3%. Repeat elements represented 63.8% of the genome, with LTR elements being the most abundant. DRAGO3 predicted 2,310 disease resistance-like genes across the 12 chromosomes of SB22; the MEME suite was used to identify their amino acid motifs. Putative candidate genes contributing to CRKN resistance were mapped on chromosome 11 of SB22. The SB22 draft genome is a valuable genomic resource for potato breeding programs.

## Introduction

Potato *(Solanum tuberosum),* a tuber crop originating in the Andes mountains of South America, is the world’s fourth most important food crop after rice, wheat, and maize. It is a new world crop introduced in the late sixteenth century and spread worldwide by European settlers. The first potato patch in the United States of America (US) was planted in 1719 in New Hampshire (Wilson, 1959). With 410 million cwt valued at $3.9 billion in 2021 (USDA, 2022), potato ranks first among the most produced vegetable crops in the US. Potatoes are processed (french fries, chips, canned and dehydrated products) and sold for fresh consumption, seed material, and livestock feed. The Pacific Northwest, which includes Oregon, Washington, and Idaho, is a highly productive potato-growing region, accounting for ∼60% of global production. The value of potatoes sold by these three states was $1.92 billion in 2021 (Potato Grower, 2022). The Tri-State potato research and breeding program is a regional collaboration among the state universities of Idaho, Oregon, and Washington. It was formed in 1985 to develop potato varieties that meet the growing demands of the tri-state region. The Tri-State breeding program aims to develop potatoes with superior internal and external qualities, early maturity and long shelf-life, high yield, and biotic and abiotic stress tolerance. As of 2021, the program has released 39 potato varieties cultivated in the tri-state region and other parts of the US (PVMI, 2023). The program aims to incorporate advanced genomic resources and marker-assisted selection approaches to accelerate the breeding process, which typically takes 10-12 years before new cultivar release, and to discover new biotic and abiotic stress tolerance/resistance sources in available germplasm.

Potatoes are susceptible to various diseases caused by viruses, fungi, bacteria, and nematodes. In the tri-state region, the root-knot disease caused by Columbia root-knot nematode (CRKN) (*Meloidogyne chitwoodi Golden* O’Bannon, Santo, and Finley) is a significant pest having a severe economic impact on the potato crop (King and Taberna 2013). The nematode was first described in 1980 and is known to infect such crops as potatoes, wheat, corn, and barley (Santo 1980). It is an endo-parasitic nematode that establishes itself by feeding on potato tubers, causing galls or blemishes on the potato surface and internal defects (Elling 2013). Potatoes with visual defects covering >5 % of the tuber surface area are rejected for fresh market, and >5-15 % are rejected by processors, making the crop commercially nonviable (Ingham *et al*. 2007). CRKN spreads primarily through seed materials, irrigation water, and soil on farm equipment. The nematode also has a wide variety of hosts in monocot and dicot plants, including cultivated crops and weeds, making it difficult to eradicate (Santo *et al*. 1988; Ferris *et al*. 1993; Rich *et al*. 2009). Standard control measures include crop rotation, nematicide applications, and planting resistant varieties. CRKN-resistant varieties provide the most cost-effective and environmentally sustainable approach to CKRN control. *Solanum tuberosum* lacks the natural resistance to *M. chitwoodi* found in closely related wild S*olanum* species *S. bulbocastanum, S. cardiophyllum, S. brachistotrichum, S. Fendleri,* and *S. hougasii*. (Janssen *et al*. 1995).

*Solanum bulbocastanum* Dunal is an ornamental night shade in the family *Solanaceae*. It is a wild diploid species (2*n* = 2*x* = 24) originating from Mexico, with an endosperm balance number (EBN) of 1. Sasser et al. (1984) showed that *S. bulbocastanum* exhibited high resistance to root-knot nematodes. Accessions of *S. bulbocastanum* are also known to be moderately resistant to *Meloidogyne persicae* and *Meloidogyne euphorbiae* (Radcliffe and Lauer, 1966; Radcliffe *et al*. 1981). Brown (1989) indicated that *S. bulbocastanum* has significant resistance to *M. chitwoodi* race 1 and race 2*. S. bulbocastanum* was a source used to impart natural resistance to CRKN in commercial potato cultivars through somatic hybridization (Austin *et al*. 1993) and subsequent introgression of resistance to *S. tuberosum* through backcrossing (Brown *et al*. 1995, 1999). The gene *R_mc1_* that confers resistance to race 1 of *M. chitwoodi* was mapped to chromosome 11 of SB22 (Brown *et al*. 1996) and QTLs associated with *M. chitwoodi* resistance were identified from resistant lines (Zhang *et al*. 2007). Resistance to root-knot disease is conferred by two known *R* genes, *RMctuber(blb)* and *RMc(blb1)* (Brown *et al*. 2009). SB22 is the parent species of the CRKN (race 1 and race 2) resistant genotype PA99N82-4 developed by somatic hybridization of SB22 with *S. tuberosum* and further backcrossing of hybrids with *S. tuberosum* (Brown *et al*. 2009; Graebner *et al*. 2018). Studies on resistant and susceptible potato varieties indicated the inability of CRKN juveniles to establish feeding sites and grow in resistant varieties (Brown *et al*. 2014; Bali *et al*. 2019). Transcriptomic profiling of susceptible and *S. bulbocastanum* introgressed resistant lines indicated upregulation of defense-related genes (Bali *et al*. 2019). Although CRKN is an important parasite in the tri-state region and a resistance gene candidate has been identified, knowledge about molecular mechanisms surrounding resistance is limited.

Identification of plant disease resistance (“*R”*) genes and associated markers will be valuable to potato breeding programs. Most *R* genes encode proteins that contain a putative amino-terminal signaling domain, a nucleotide binding site (NBS), and a series of carboxy-terminal leucine-rich repeats (LRR). These *R* genes are further divided into TIR-NBS-LRR (TNL), which has an amino-terminal TIR (Toll/interleukin receptor), and CC-NBS-LRR (CNL), which encodes an amino-terminal coiled-coiled motif (van Wersch *et al*. 2020; Li *et al*. 2021; Wu *et al*. 2021).

*R* genes obtained from *S. bulbocastanum* are an essential source of resistance to late blight, and exhibit long-term durable resistance to all races of *Phytophthora infestans* (Helgeson *et al*. 1998). A major *R*-gene locus (RB) conferring resistance to late blight between markers CP53 and CT64 was mapped to chromosome 8 with map-based approaches (Naess *et al*. 2000). Transgenic lines containing *R* genes belonging to the CNL (coiled coil-nucleotide binding site-leucine-rich repeat) class exhibited resistance to late blight (Song *et al*. 2003). Similarly, other *R*-genes like *Rpi-blb2* and *Rpi-blb3* from *S. bulbocastanum* were mapped and introgressed into potato germplasm to provide resistance to *P. infestans* (Park *et al*. 2005; Orbegozo *et al*. 2016; Sanetomo *et al*. 2019; Rakosy-Tican *et al*. 2020).

The CRKN-resistant line of *S. bulbocastanum*, PI 275187, selection no. 22 (SB22), was collected from Michoacán de Ocampo, Mexico. SB22 was found to be highly resistant to *M. chitwoodi* races 1 and 2, as well as high regeneration efficiency in protoplasmic fusion with *S. tuberosum* (Brown 1989; Austin *et al*. 1993). Given the importance of *S. bulbocastanum* as a source of disease and pest resistance, the SB22 genome sequencing project was undertaken to expand knowledge of the *S. bulbocastanum* genome to better understand current resistance mechanisms and to address the inevitable emergence of new races of pests and pathogens in the future.

## Methods and Materials

### Plant material

SB22 (PI 275187) was obtained as a clonal selection from USDA, National Plant Germplasm Systems by Dr Charles Brown (USDA-ARS, Prosser, WA). High molecular weight genomic DNA was isolated from 1-3g of young, unexpanded leaf tissue using the method of Healey et al. (2014), with the final gDNA sample re-suspended in EB buffer (Qiagen, Hilden, Germany). Genomic DNA quality was assessed using a Nanodrop spectrophotometer with accepted 260/280 absorbance ratio values greater than or equal to 1.8 and by gel electrophoresis of both intact and HindIII-digested samples to ensure integrity and absence of contaminating RNA and polysaccharides. DNA quantity was measured by Qubit, using the Qubit® dsDNA HS Assay Kit (Molecular Probes, Eugene, OR). DNA was sequenced on an Pacific Biosciences RSII instrument at the Arizona Genomics Institute. Sequencing libraries were prepared at 20 and 35 kb and loaded on 44 SMRT cells.

### Genome assembly

An initial genome assembly was generated from the PacBio data using Falcon v0.5 (Chin *et al*. 2016). Errors were corrected using Illumina HiSeq3000 reads (2×150 bp) with Pilon v1.22 (Walker *et al*. 2014). Genome scaffolding using the HiRise method was performed by Dovetail Genomics (Scotts Valley, CA, USA). BUSCO (Manni *et al*. 2021) was invoked to assess genome completeness using the Solanaceae_odb10 data set (Manni *et al*. 2021). Heterozygosity and genome size were estimated using a Kmer-based approach (k = 21) with Illumina reads, using Jellyfish version 2.3 (Marçais and Kingsford 2011) and GenomeScope (Vurture *et al*. 2017). As another check for completeness, the LTR assembly index (LAI) score was computed using the tools ltr_harvest version v1.6.1 (Ellinghaus *et al*. 2008), LTR_FINDER_parallel v1.07 (Xu et al. 2007), and LTR_retriever (Ou and Jiang 2018).

### Synteny analysis

The Genome assembly of SB22 was aligned to DM6.1 potato genome (Pham et al. 2020) with minimap2 v2.26 (Li 2018). The alignment was processed through Syri v1.6.3 (Goel *et al*. 2019) to predict the structural variations and the final predictions were plotted using Plotsr v1.1.0 (Goel and Schneeberger 2022).

### Repeat masking and functional annotation

To model repeats in the genome, de-novo repeat analysis was carried out using the RepeatModeler v2.0.4 package (Flynn et al. 2020). The repeat library generated from RepeatModeler was used to mask the repeat regions using RepeatMasker v4.1 (Nishimura 2000) and to estimate the total repeat content in the SB22 genome. The repeat masked genome was used in BRAKER pipeline version 2.1.6 (Stanke *et al*. 2006, 2008; Hoff *et al*. 2016, 2019; Brůna *et al*. 2021) with PA99N82–4 transcriptome (Bali *et al*. 2019) containing >30 million Illumina reads (1×150 bp) obtained from roots and a set of curated proteins from Viridiplantae-ODB10, and gene models from the DM6 potato genome (Pham et al.2020). BUSCO protein was analyzed using the finalized predicted transcripts and Solanales_odb10. Identified gene models were annotated using the KEGG database for pathway analysis, and functional annotation was performed using EGGNOG-mapper web v2.0.1 (Huerta-Cepas *et al*. 2019; Cantalapiedra *et al*. 2021) against the eggNOG 5 database. Identified gene models were comparatively analyzed using TRAPID tool with EggNOG 4.5 database as the reference (Van Bel *et al*. 2013). Microsatellite analysis for the entire SB22 genome was carried out using MISA v2.1 (Thiel *et al*. 2003) microsatellite analysis tool using default parameters (minimum 7/di, 5/tri, 5/tetra, 5/penta and 5/hexa nucleotide repeats).

### R gene Prediction and analysis

Predicted gene models from SB22 were subject to *R*-gene motif identification using DRAGO3 (Osuna-Cruz et al. 2017), which uses HMMER v3 package and 60 HMM modules along with CC and TM domains using COILS 2.2 and TMHMM 2.0c programs. The predicted *R*-genes were further subjected to motif prediction using the MEME Suite (Bailey *et al*. 2015) with the number of motifs to predict set to 10. In addition, CRKN resistance markers previously reported by (Bali *et al*. 2022) were mapped to the assembled SB22 genome to identify putative candidate genes.

## Results and Discussion

### Genome Assembly

PacBio RSII sequencing yielded 39 Gb, an estimated genome coverage of 36x. Genome size estimation by Jellyfish and GenomeScope gave an estimated haploid genome size of 741 Mb, with a heterozygosity of ∼1.85% (Figure S1). Following Falcon assembly and error correction with Illumina reads using Pilon, an initial assembly consisting of 655,247,856 bp was produced which had 1,301 contigs, ranging in size from 29,502,347 bp to 805 bp (Table 1). The Dovetail HiRise assembly significantly increased contiguity, reducing the number of genome scaffolds to 406 (NCBI project id: PRJNA1003451) and increasing the total assembly size to 656,373,245 bp and N50=2,211.9 kb. The average length of the scaffolds was 1,616,683 bp, and the maximum length was 60,441,652 bp. The genome assembly had a high degree of completion for Solanaceae with a reported BUSCO score of 95.7 % (Table 2).

**Table 1:** Assembly metrics of *S. bulbocastanum*.

|  |  |
| --- | --- |
| Main genome scaffold total: | 406 |
| Main genome contig total: | 1313 |
| Main genome scaffold sequence total: | 656373 Kb |
| Main genome contig sequence total: | 656277 Kb (0.015% gap) |
| Main genome scaffold N/L50: | 6/51167 Kb |
| Main genome contig N/L50: | 77/2212 Kb |
| Main genome scaffold N/L90: | 13/24490 Kb |
| Main genome contig N/L90: | 413/248573 Kb |
| Max scaffold length: | 60442 Kb |
| Max contig length: | 29550 Kb |
| Number of scaffolds > 50 KB: | 165 |
| % main genome in scaffolds > 50 KB: | 99.13% |

**Table 2:** BUSCO analysis of SB22 genome against Solanaceae-odb10 database.

| No of Busco's | Completeness |
| --- | --- |
| 5,693 (95.7%) | Complete BUSCOs (C) |
| 5,367 (90.2 %) | Complete and single-copy BUSCOs (S) |
| 326 (5.5 %) | Complete and duplicated BUSCOs (D) |
| 65 (1.1%) | Fragmented BUSCOs (F) |
| 192 (3.2 %) | Missing BUSCOs (M) |
| 5,950 | Total BUSCO groups searched |

The SB22 genome assembly contained more chromosomal inversions than translocations relative to the *Solanum tuberosum* reference, DM6. Chromosome 11, which includes markers for CRKN resistance, showed two small inversions near the centromeric region, while both arms of the chromosome showed high synteny to DM6.1 (Figure 1). This pattern of pericentromeric inversions was seen on most of the chromosomes.

**Figure 1:**
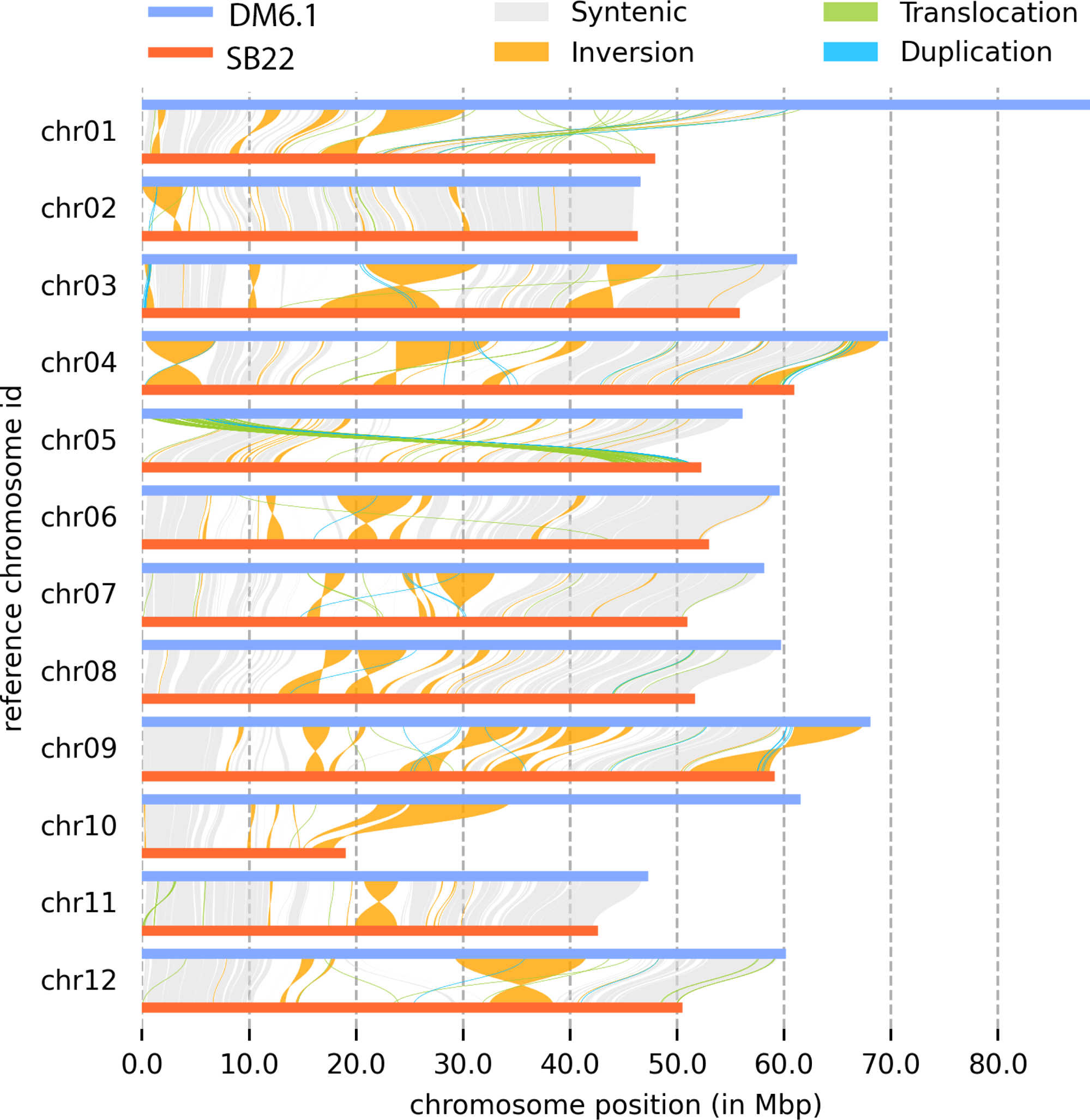
Syri plot constructed using alignment between DM6.1 genome and Pseudomolecules of SB22,

### Repeat Analysis

De-novo repeat analysis with RepeatModeler and RepeatMasker estimated a repeat content of ∼419 Mb, representing 63.83 % of the total assembled genome (Table 3). Retroelements represented 31.63 % of the genome with 1,85,113 repeat elements (Figure 2). LTR elements were the second most abundant, representing 29.27 % with 29,738 elements, followed by Gpysy/DIRS1 with 22.30% with 1,00,741 repeat elements.

**Figure 2:**
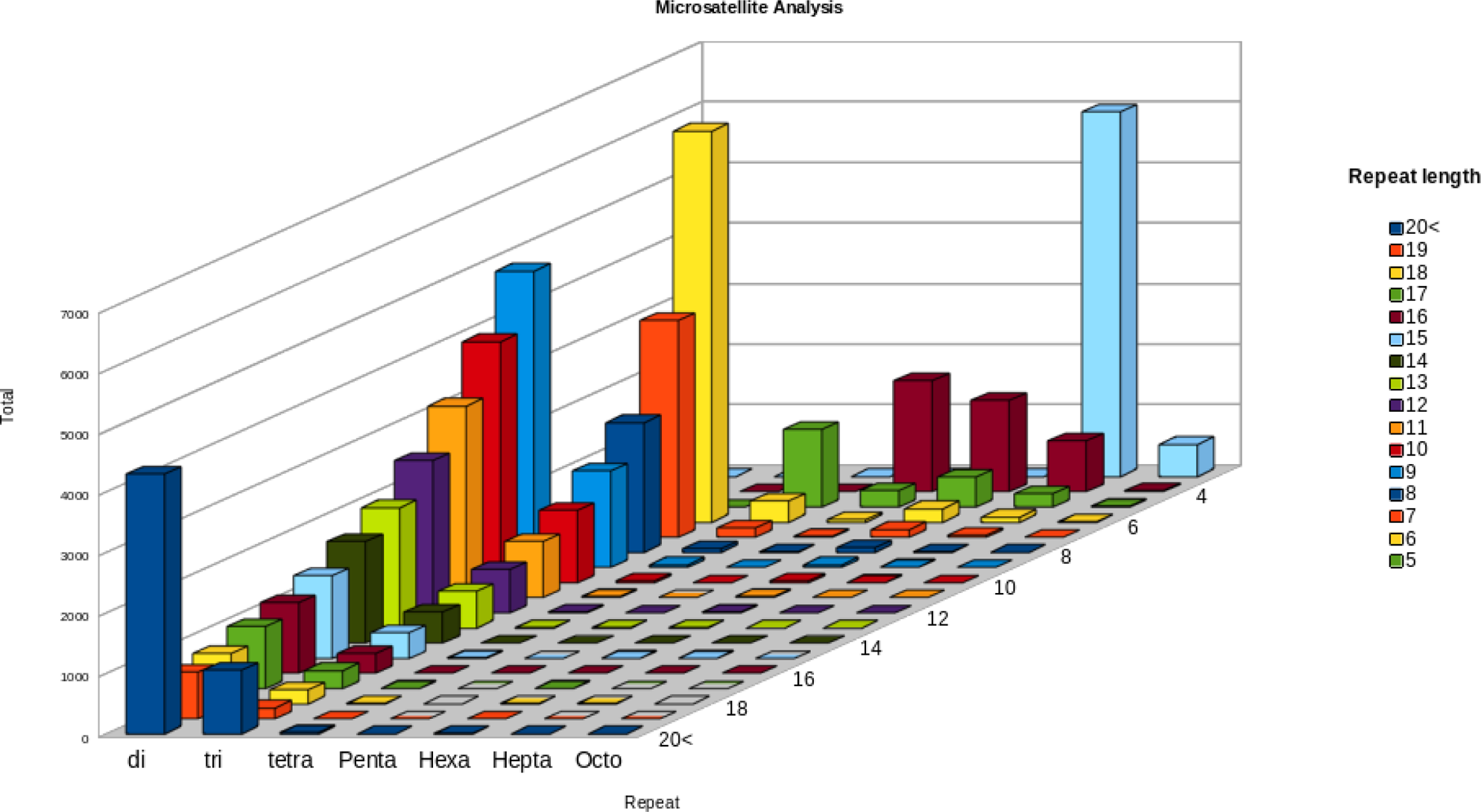
Genomic Features of SB22 Genome. From the outer circle gene density, repeat positions and positions of CNL and TNL genes are identified in the chromosomes. A - Gene Models, B - LTR-Gypsy, C - LTR-Copia, D - LINE-RTE-BovB, E - LINE-L1, F - Simple_repeats, G – unknown repeats

**Table 3:**
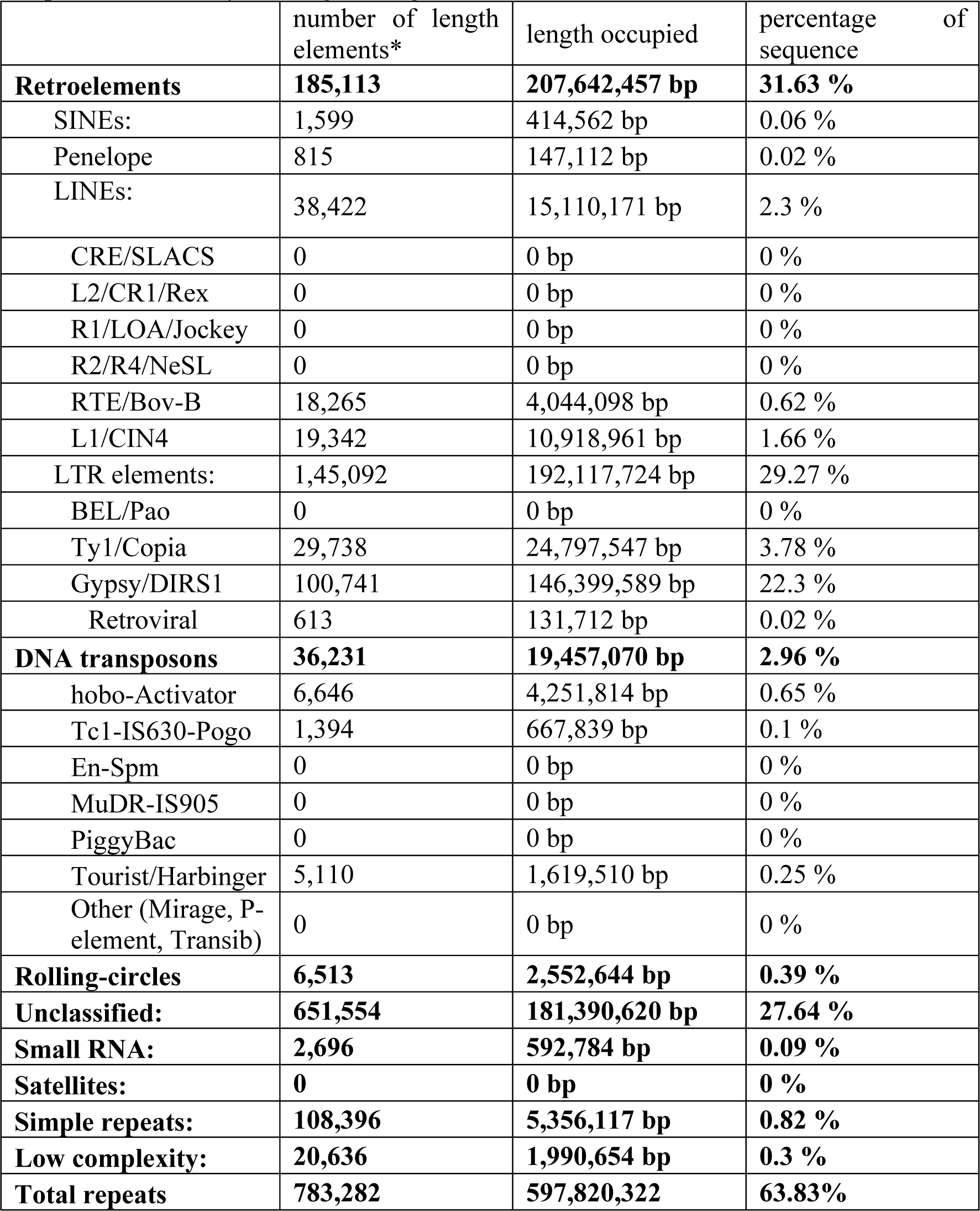
Table showing the abundance of various repeat categories predicted by Repeat modeler and Repeat masker analysis using SB22 genome.

|  | number of length<br>elements* | length occupied | percentage of<br>sequence |
| --- | --- | --- | --- |
| **Retroelements** | **185,113** | **207,642,457 bp** | **31.63 %** |
| SINEs: | 1,599 | 414,562 bp | 0.06 % |
| Penelope | 815 | 147,112 bp | 0.02 % |
| LINES: | 38,422 | 15,110,171 bp | 2.3 % |
| CRE/SLACS | 0 | 0 bp | 0 % |
| L2/CR1/Rex | 0 | 0 bp | 0 % |
| R1/LOA/Jockey | 0 | 0 bp | 0 % |
| R2/R4/NeSL | 0 | 0 bp | 0 % |
| RTE/Bov-B | 18,265 | 4,044,098 bp | 0.62 % |
| L1/CIN4 | 19,342 | 10,918,961 bp | 1.66 % |
| LTR elements: | 1,45,092 | 192,117,724 bp | 29.27 % |
| BEL/Pao | 0 | 0 bp | 0 % |
| Ty1/Copia | 29,738 | 24,797,547 bp | 3.78 % |
| Gypsy/DIRS1 | 100,741 | 146,399,589 bp | 22.3 % |
| Retroviral | 613 | 131,712 bp | 0.02 % |
| **DNA transposons** | **36,231** | **19,457,070 bp** | **2.96 %** |
| hobo-Activator | 6,646 | 4,251,814 bp | 0.65 % |
| Tc1-IS630-Pogo | 1,394 | 667,839 bp | 0.1 % |
| En-Spm | 0 | 0 bp | 0 % |
| MuDR-IS905 | 0 | 0 bp | 0 % |
| PiggyBac | 0 | 0 bp | 0 % |
| Tourist/Harbinger | 5,110 | 1,619,510 bp | 0.25 % |
| Other (Mirage, P-element, Transib) | 0 | 0 bp | 0 % |
| **Rolling-circles** | **6,513** | **2,552,644 bp** | **0.39 %** |
| **Unclassified:** | **651,554** | **181,390,620 bp** | **27.64 %** |
| **Small RNA:** | **2,696** | **592,784 bp** | **0.09 %** |
| **Satellites:** | **0** | **0 bp** | **0 %** |
| **Simple repeats:** | **108,396** | **5,356,117 bp** | **0.82 %** |
| **Low complexity:** | **20,636** | **1,990,654 bp** | **0.3 %** |
| **Total repeats** | **783,282** | **597,820,322** | **63.83%** |

Microsatellite prediction using MISA identified 62,429 microsatellites, with 18,612 in compound formation. The relative abundance of microsatellites was 95.2 per Mb in the SB22 genome. Dinucleotide repeats were most abundant in the genome, representing 44.2% (27,649) (Figure 3), followed by tri-nucleotide and hepta-nucleotide repeats with 32.4 % (20,248) and 11.6 % (7,241), respectively. The frequency of AT repeat motifs was the most abundant in the genome, representing 87.1 % of total di-nucleotide repeats, followed by the tri-nucleotide repeat motif AAT, which represented 60.6 %.

**Figure 3:**
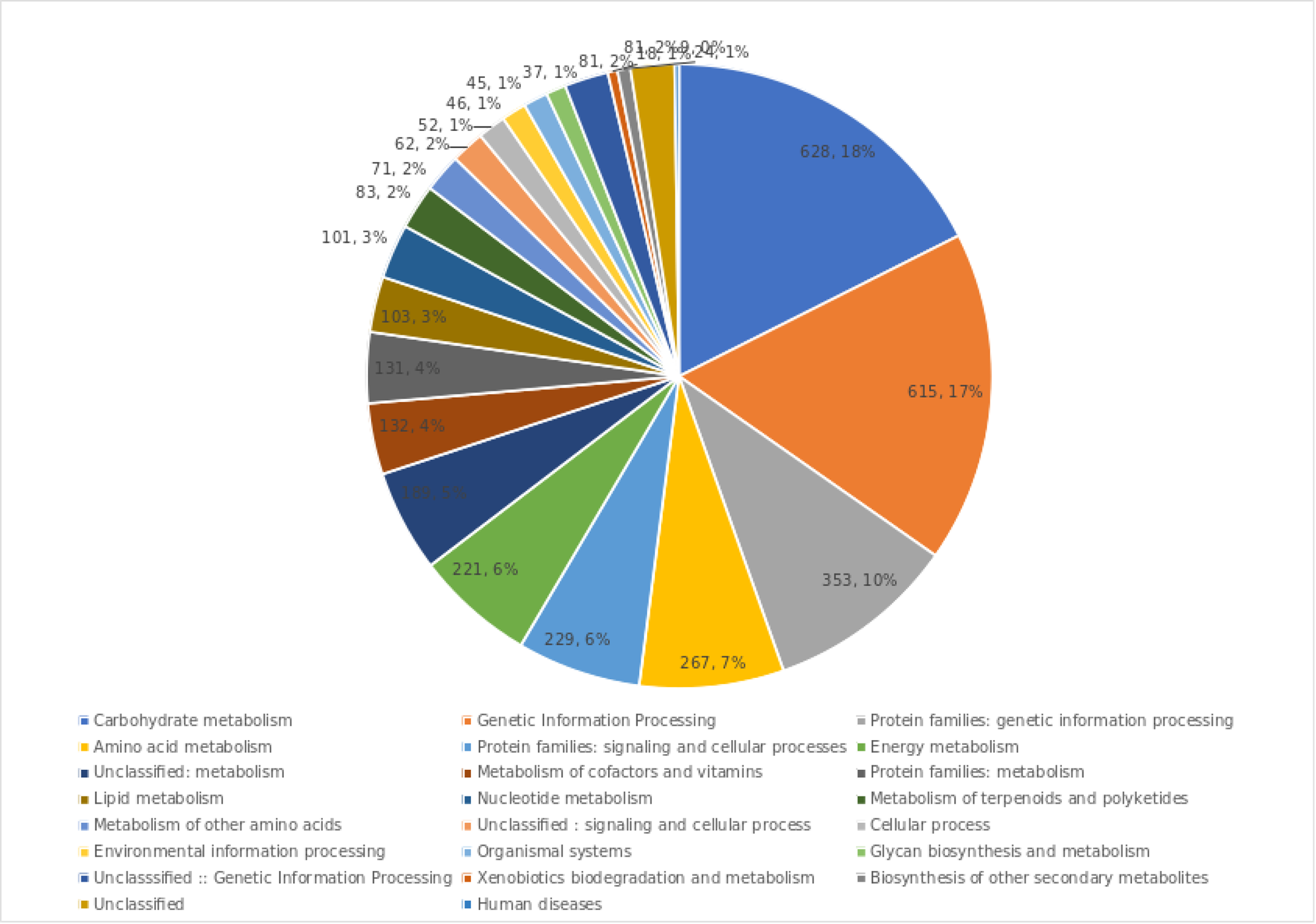
Repeat motif, length, and total number of microsatellites identified in the SB22 genome.

### Structural and Functional Annotation

The BRAKER3 gene prediction identified 43,280 transcripts. BUSCO protein analysis indicated completeness of 90.3% (Solanales_odb10) with 80.1% (3,480) complete single copy BUSCOs, 10.2% (1,197) complete duplicated BUSCOs, 3.5% (174) fragmented BUSCOs and 6.2 % (1,099) missing BUSCOs. The low completeness scores in predicted gene models may be attributed to the RNA evidence used for the functional annotation; more appropriate RNA evidence from SB22 plant tissues is necessary to further improve completeness of the predicted gene models.

Annotation with TRAPID indicated an average transcript length of 977.4 bp with 77.4% (33,490) full-length meta-annotation, 11.3 % (5,648) partial annotation, and 9.5 % (4,091) with no annotation. Of the transcript sequences, 58.85% (23,873) were similar to *Solanum tuberosum*, and 31.92% (12,676) were similar to *Solanum lycoperiscum*. A total of 7,752 gene families were identified. KEGG analysis annotated a total of 13,529 (8.2%) transcripts (Figure 4), including genetic information processing transcripts (4,058), carbohydrate metabolism (871), and Environmental Information Processing (759).

**Figure 4:**
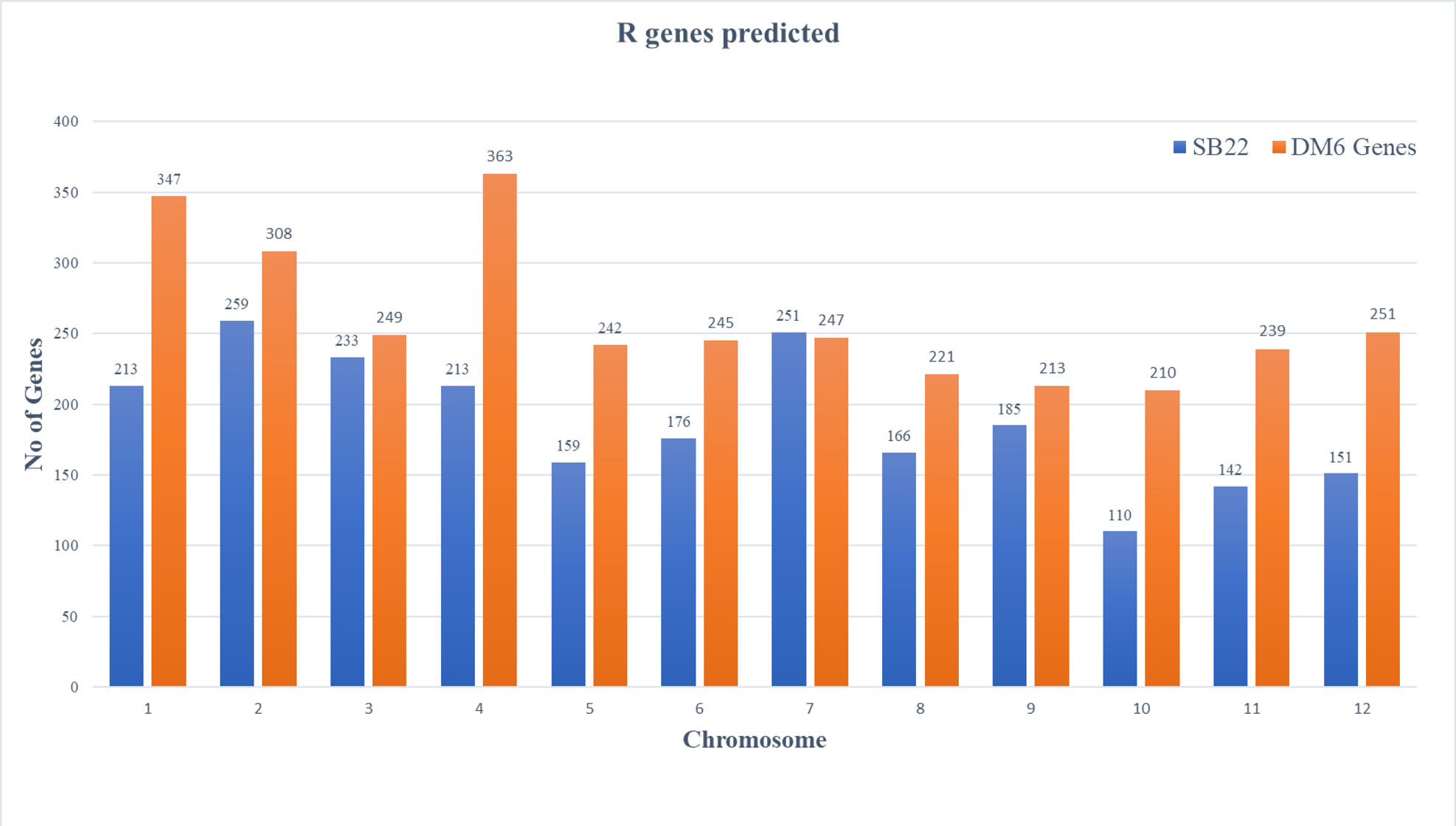
Distribution of transcripts across various gene families predicted using KEGG database

### Disease Resistance Genes

The finalized gene models generated by BRAKER3 were subject to *R*-gene prediction using DRAGO3, which identified 2,310 unique *R* genes (Table 4). *R* genes were distributed throughout the genome in SB22 across all chromosomes (Figure 5, Figure S2). DRAGO3 analysis identified genes coding for CNL, TNL, receptor-like proteins (RLP), receptor-like kinases (RLK), and other *R* gene-related domains. Among the *R* genes identified, 17 had Coiled-Coil (CC) domains, and 20 had Toll/Interleukin Receptor-like (TIR) domains (Table S1). Kinase domains were most prevalent, present in 1,441 (62 %) of the 2,310 *R* genes, followed by LRR domains, present in 789 (34 %), then NBS domains, present in 294 (12 %). Genes with two domains dominated the *R* genes identified, representing 53 % (1,233), followed by three-domain and one-domain genes with 24 % (563) and 21 % (475), respectively. Among the *R* genes, 73 candidates contained NBS-LRR domains. Annotation of the transcripts containing NBS-LRR domains using eggNOG 5 corresponded to *TMV resistance protein, TMV resistance protein N-like, Arabidopsis broad-spectrum mildew resistance protein RPW8, late blight resistance protein homolog, and Toll - interleukin 1 – resistance*, among others. The *R* gene categories containing the maximum number of domains in single gene models were CC-NBS-TM-LRR, TIR-NBS-TM-LRR, CC-Kinase-TM-LRR, and CC-TM-LECM-Kinase. CC-NBS-TM-LRR domains were the most abundant among the four domain *R* genes.

**Figure 5:**
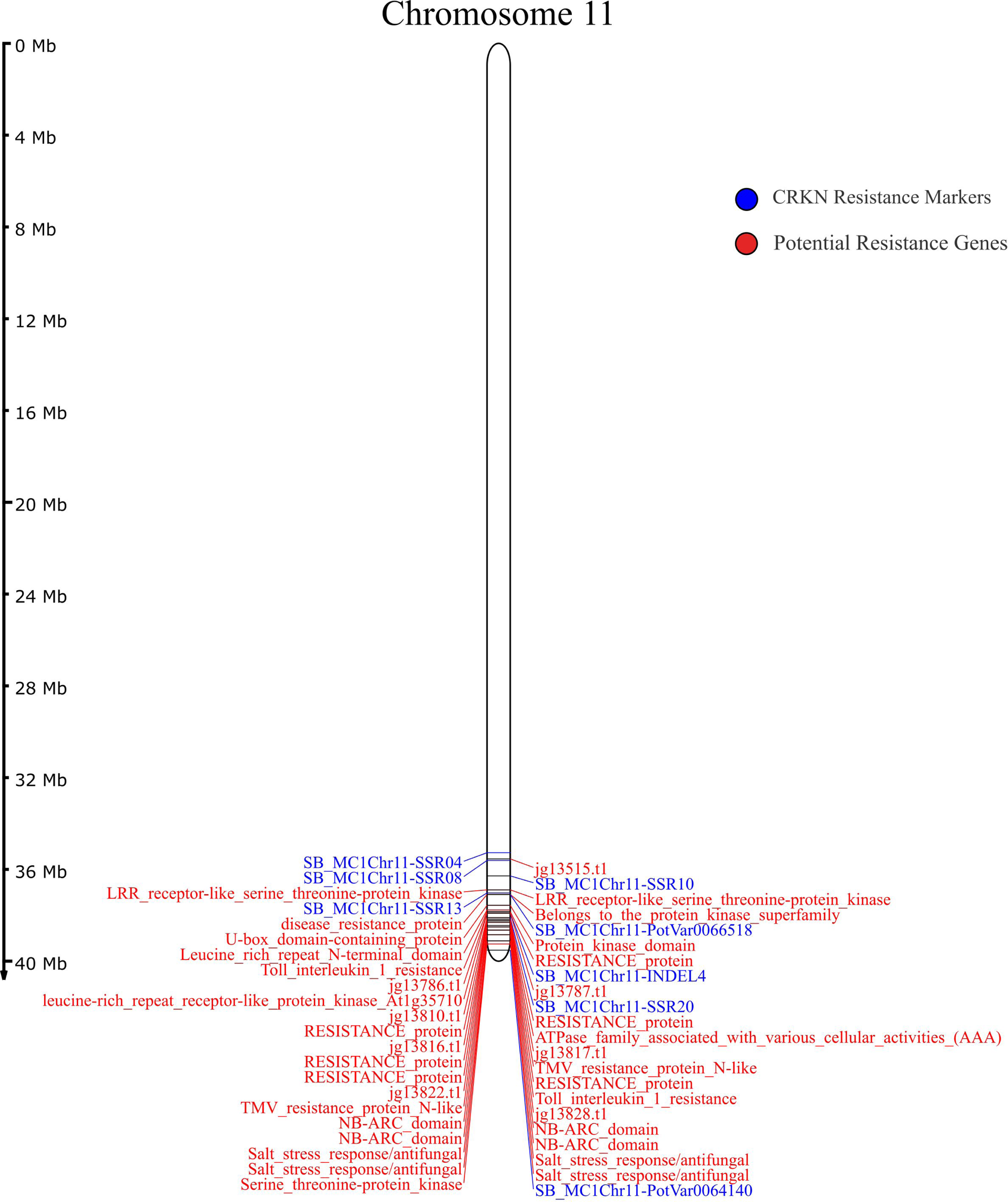
Total number of *R*-genes predicted per Chromosome of SB22 and DM6.1

**Table 4:** Comparison of numbers of R genes predicted between SB22 and DM6.1 genomes.

| R-gene | No of Genes |  |
| --- | --- | --- |
|  | SB22 | DM6.1 |
| CK | 93 | 136 |
| CL | 20 | 37 |
| CLK | 7 | 10 |
| CN | 40 | 110 |
| CNK | 0 | 0 |
| CT | 3 | 5 |
| CNL | 17 | 92 |
| KIN | 933 | 1095 |
| L | 85 | 127 |
| N | 162 | 296 |
| NK | 0 | 1 |
| NL | 52 | 121 |
| RLK | 271 | 394 |
| RLP | 304 | 388 |
| T | 18 | 25 |
| TN | 3 | 6 |
| TNL | 20 | 28 |
| TRAN | 57 | 83 |
| Total | 2310 | 3156 |

Motif prediction using MEME-identified motifs with e-values ranging from 3.2e-782 to 2.2e-755. Two motifs belonging to LRR, three protein kinases, one receptor kinase, and four motifs with unknown functions were predicted in the 2,310 R genes identified (Table 5). MEME motif prediction in CNL and TNL identified 3 LRR motifs, three coiled-coiled motifs, one P-loop, GLPL, and NBS motif (Table 6). Mapping positions of the *R* genes with CRKN resistance markers in chromosome 11 (Figure 6) of the SB22 genome located the markers near 36 *R* genes (Table: 7). Two TNL genes were identified adjacent to the markers encoding resistance gene *Toll-interleukin1-resistance,* located at 5.1 kb (upstream) and 38.5 Kb (downstream) (Figure 7) distance from the CRKN resistance markers SB_MC1Chr11-INDEL4 and SB_MC1Chr11-SSR20, evidence of their potential to contribute to CRKN resistance.

**Figure 6:**
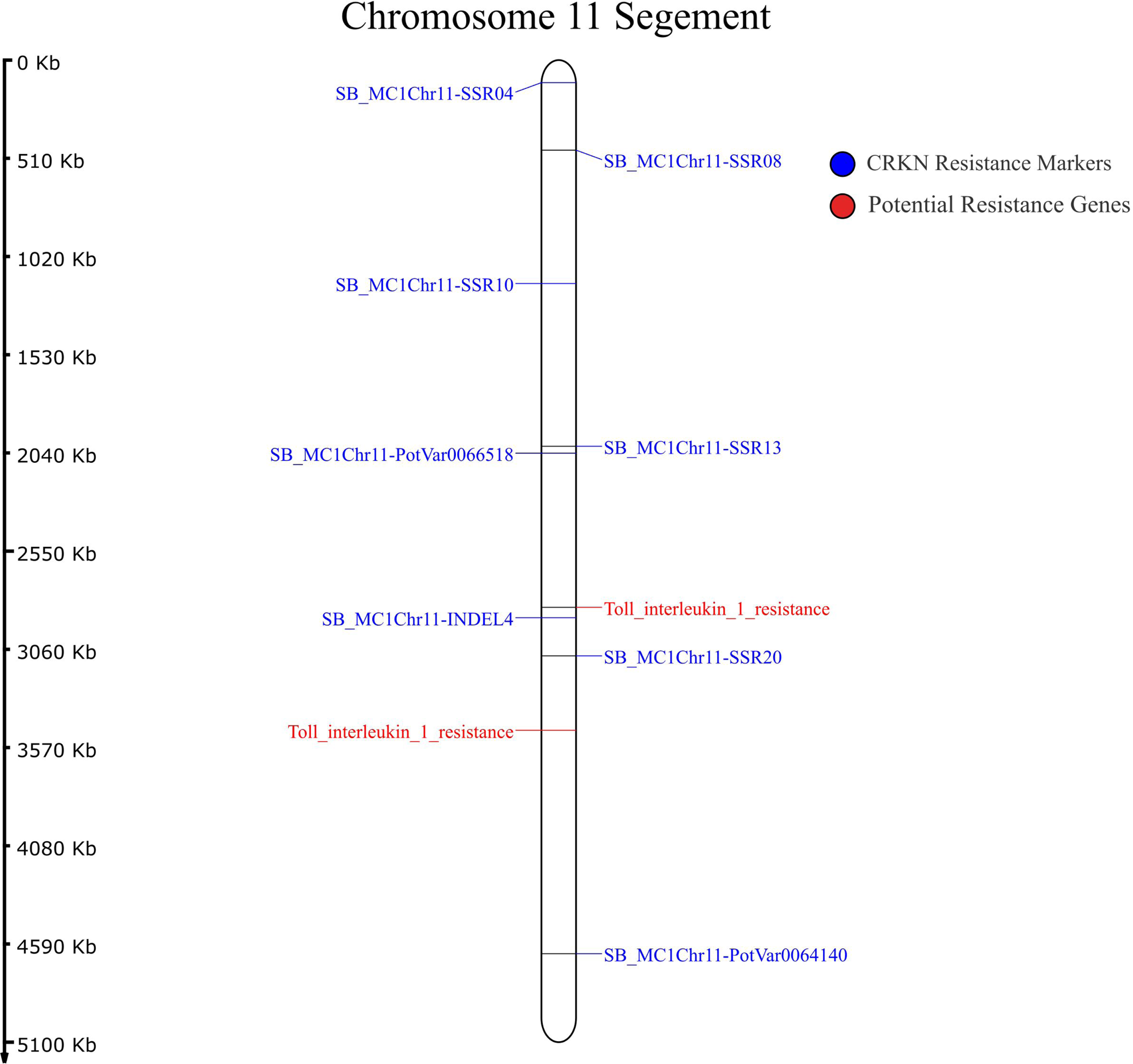
Positions of fine-mapped markers linked to CRKN resistance in Chromosome 11 of SB22

**Figure 7:**
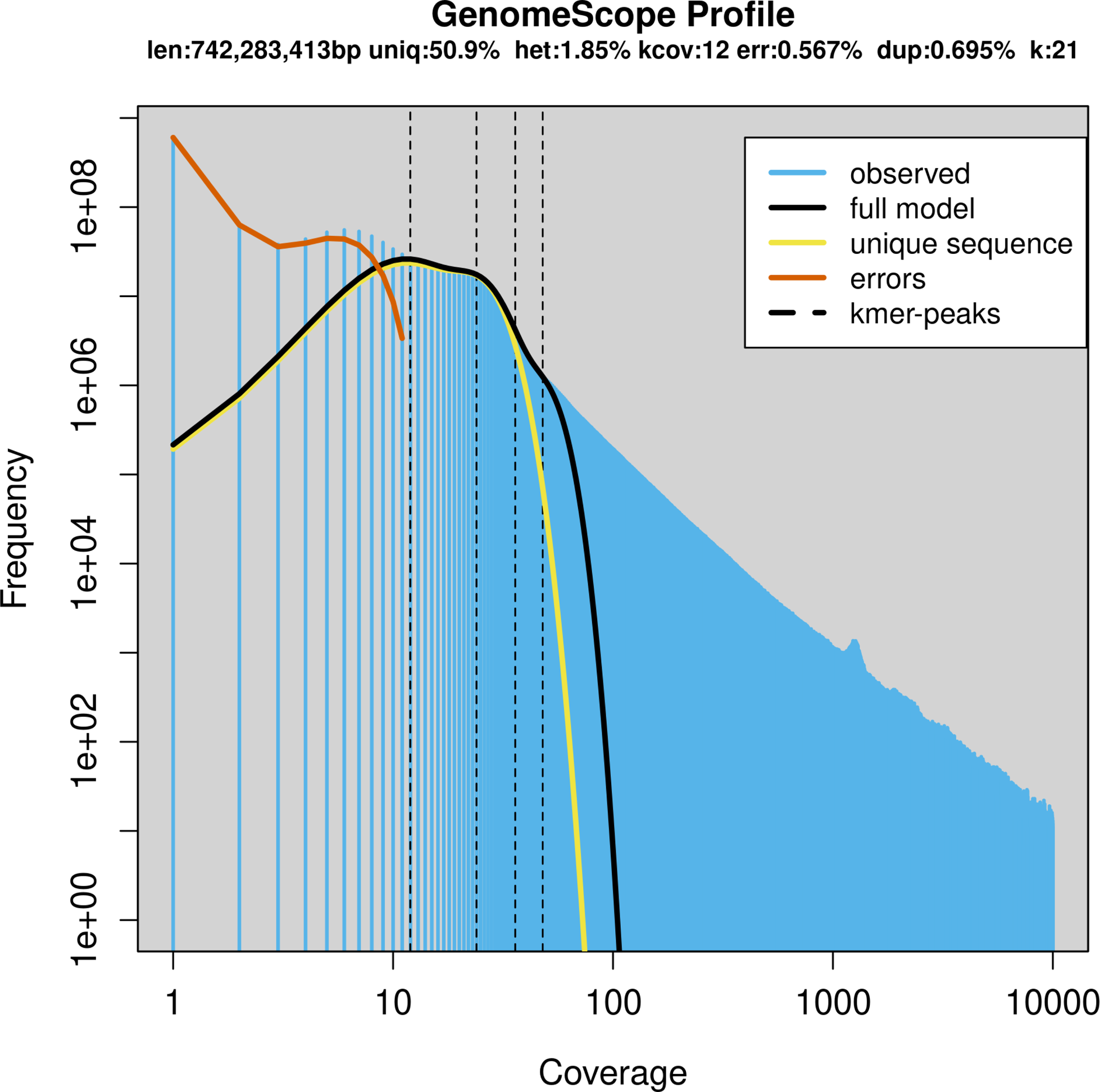
Positions of TNL genes responsible for potential CRKN resistance identified and CRKN resistance markers in the segment of chromosome 11

**Table 5:**
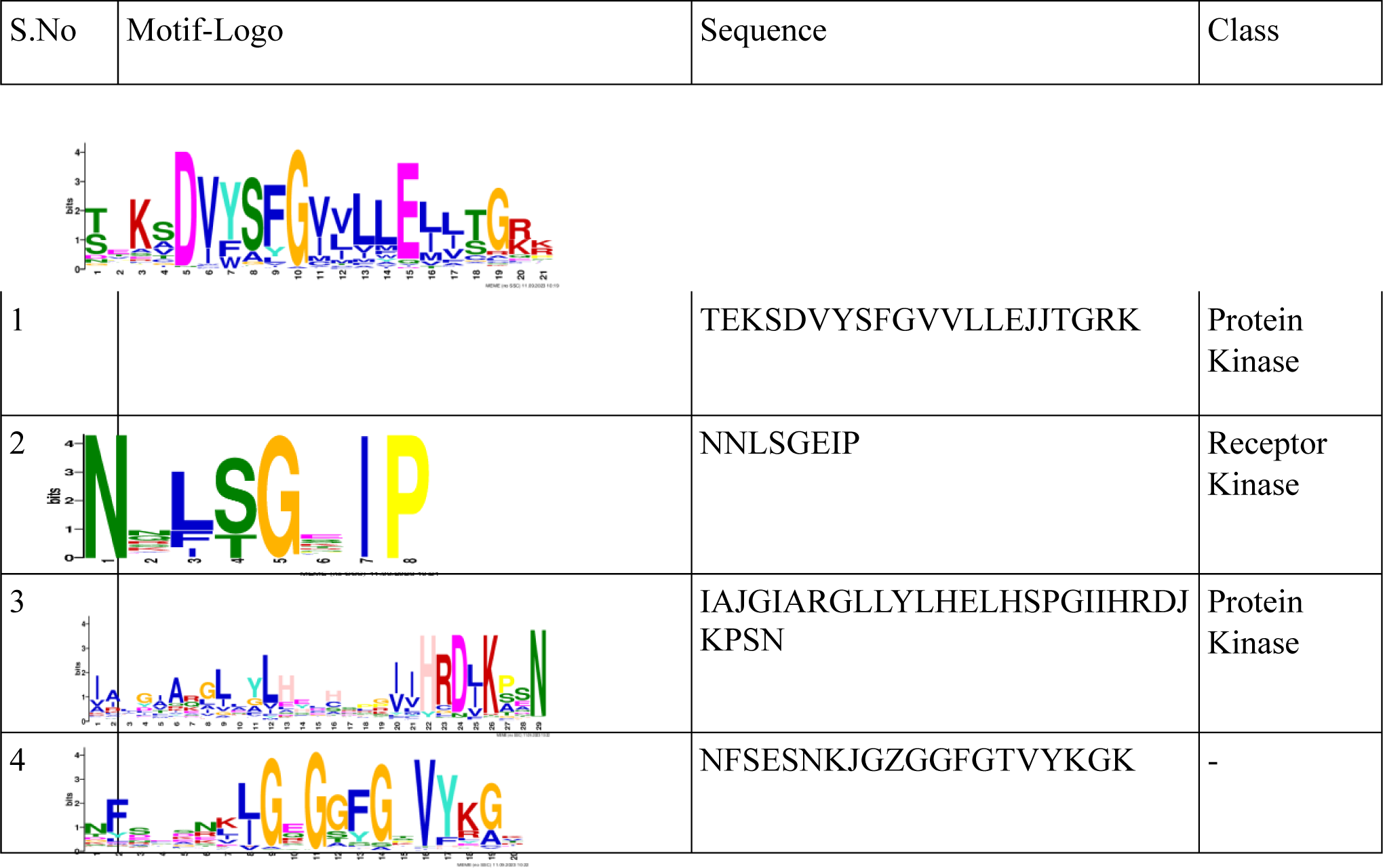
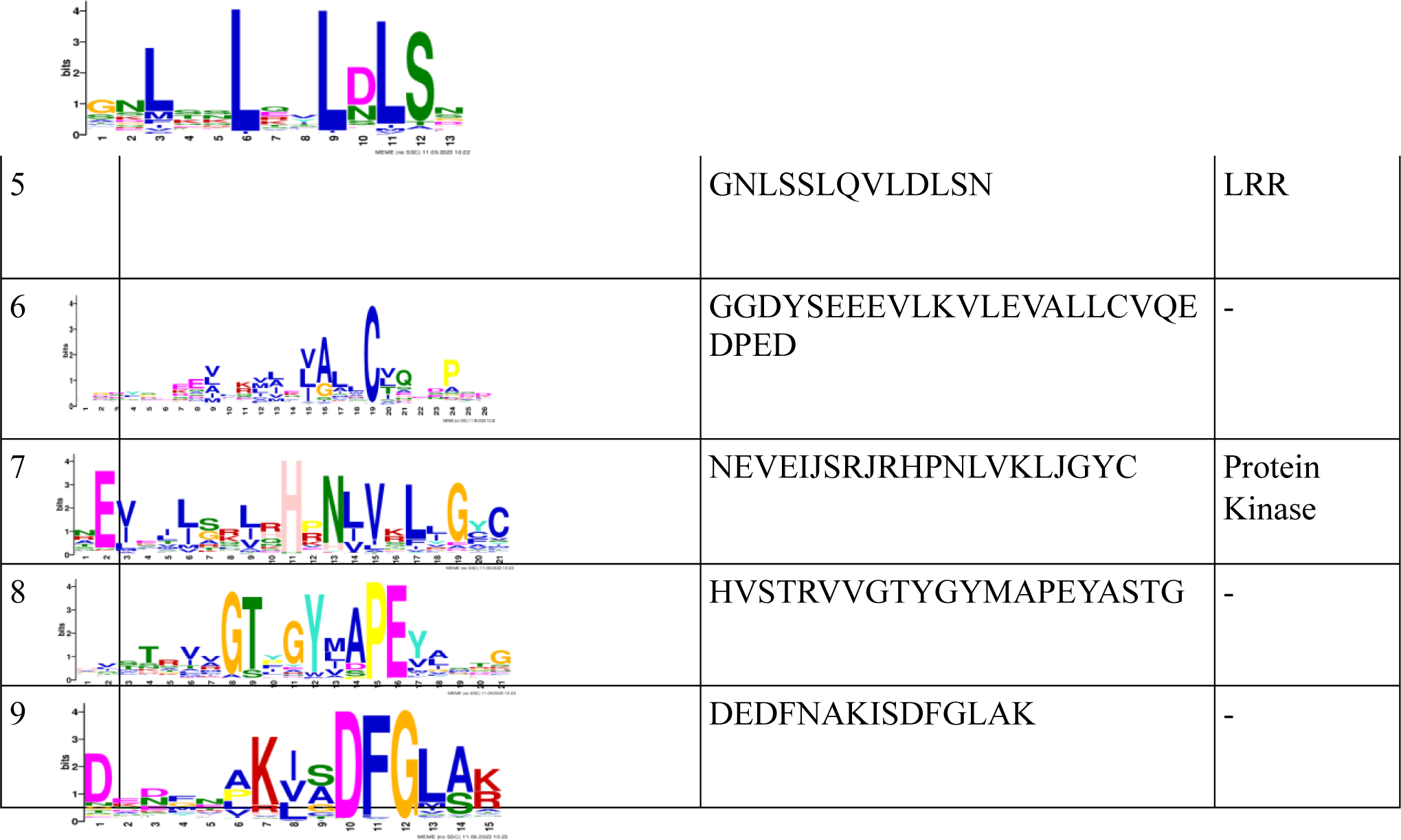
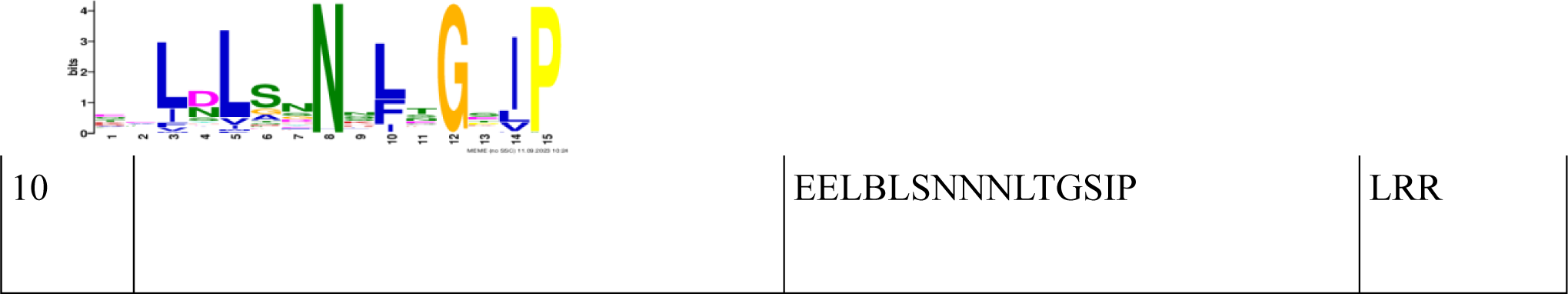
Motifs predicted by MEME tool in R genes identified in SB22 genome.

**Table 6:**
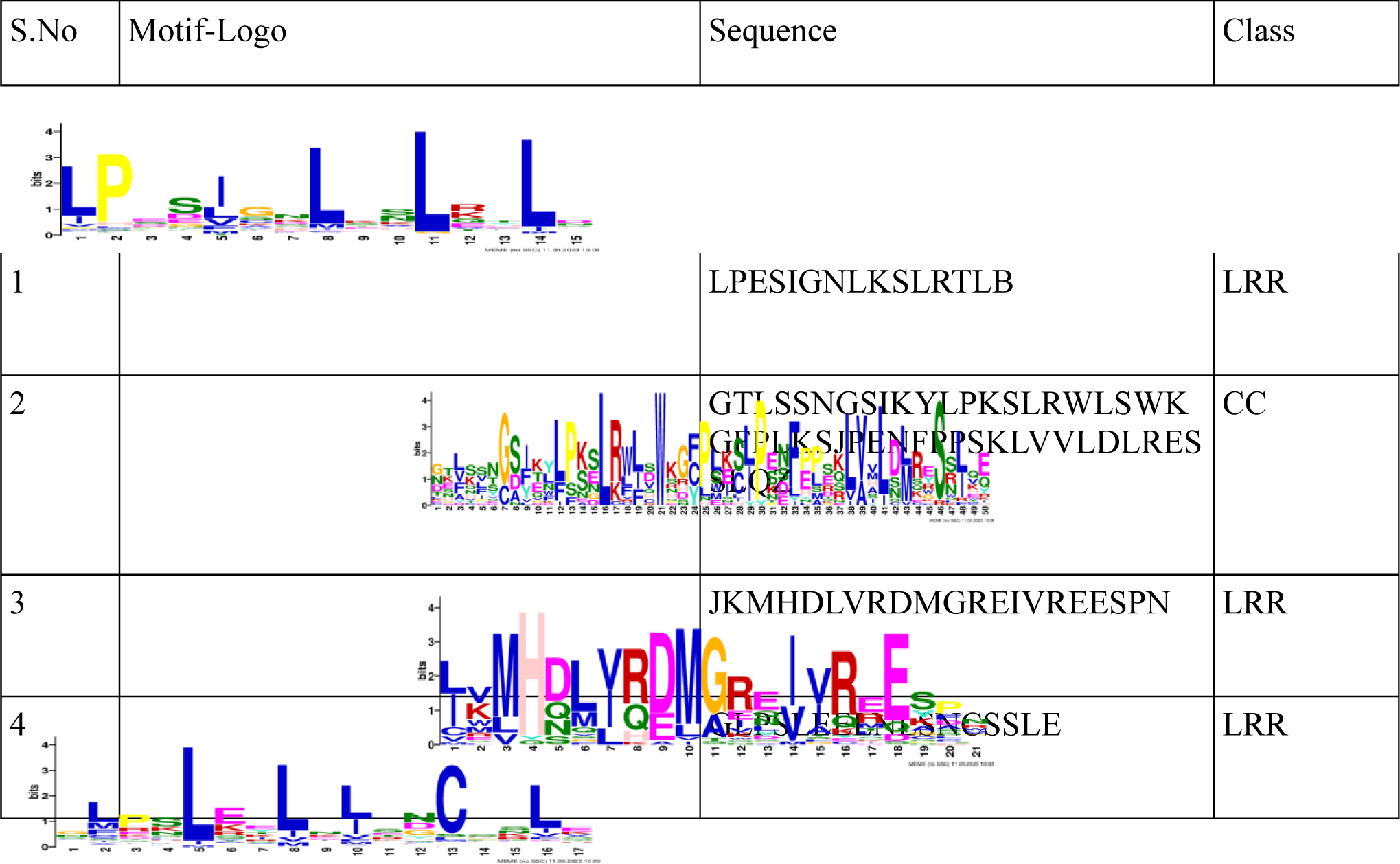
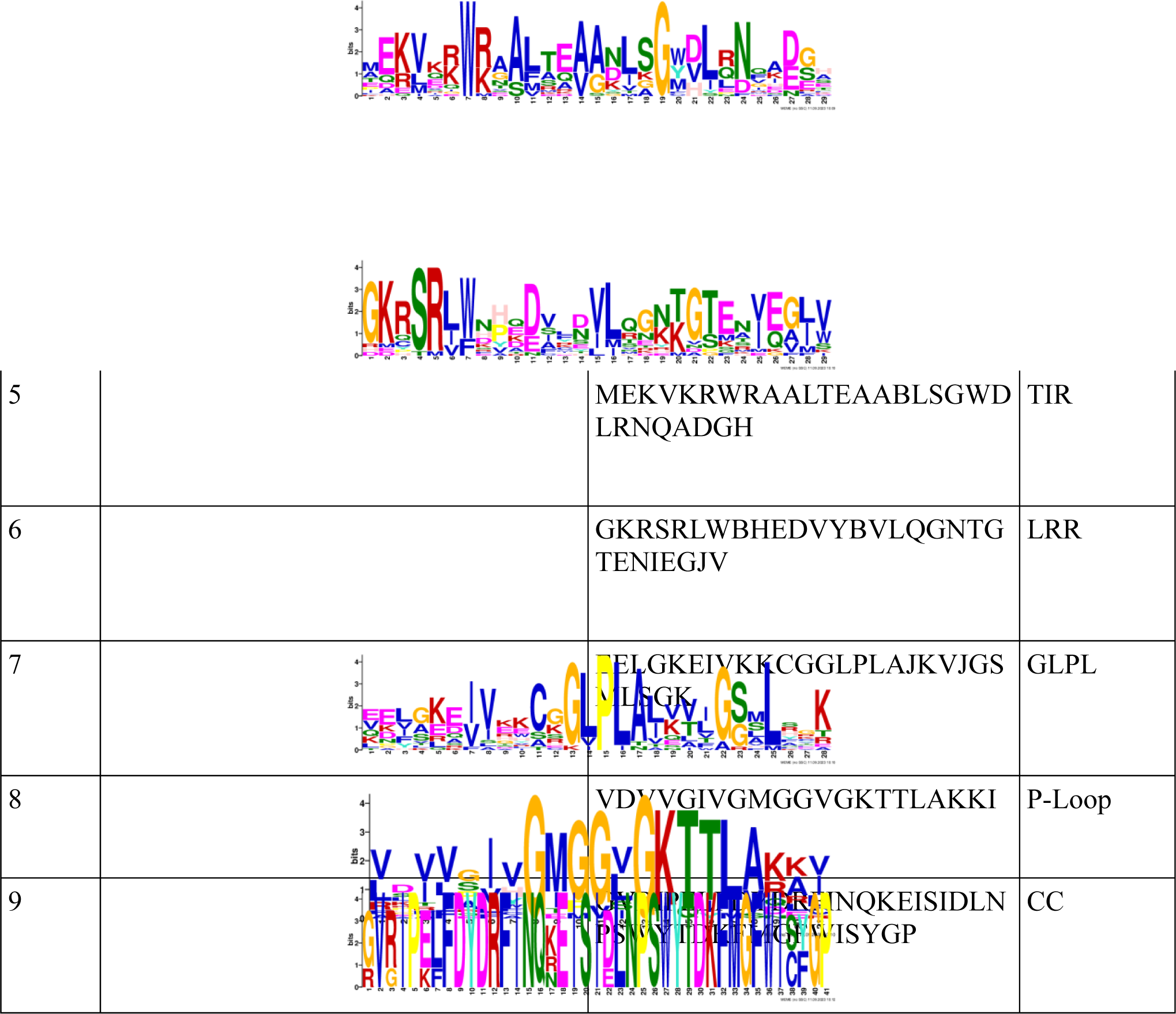
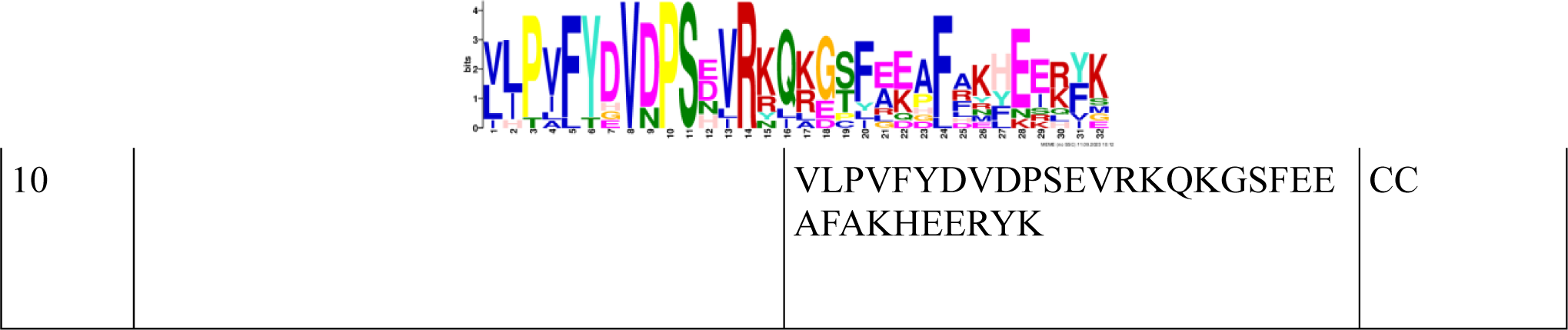
Motifs Predicted by MEME tool in CNL and TNL genes identified in SB22 genome.

**Table 7:**
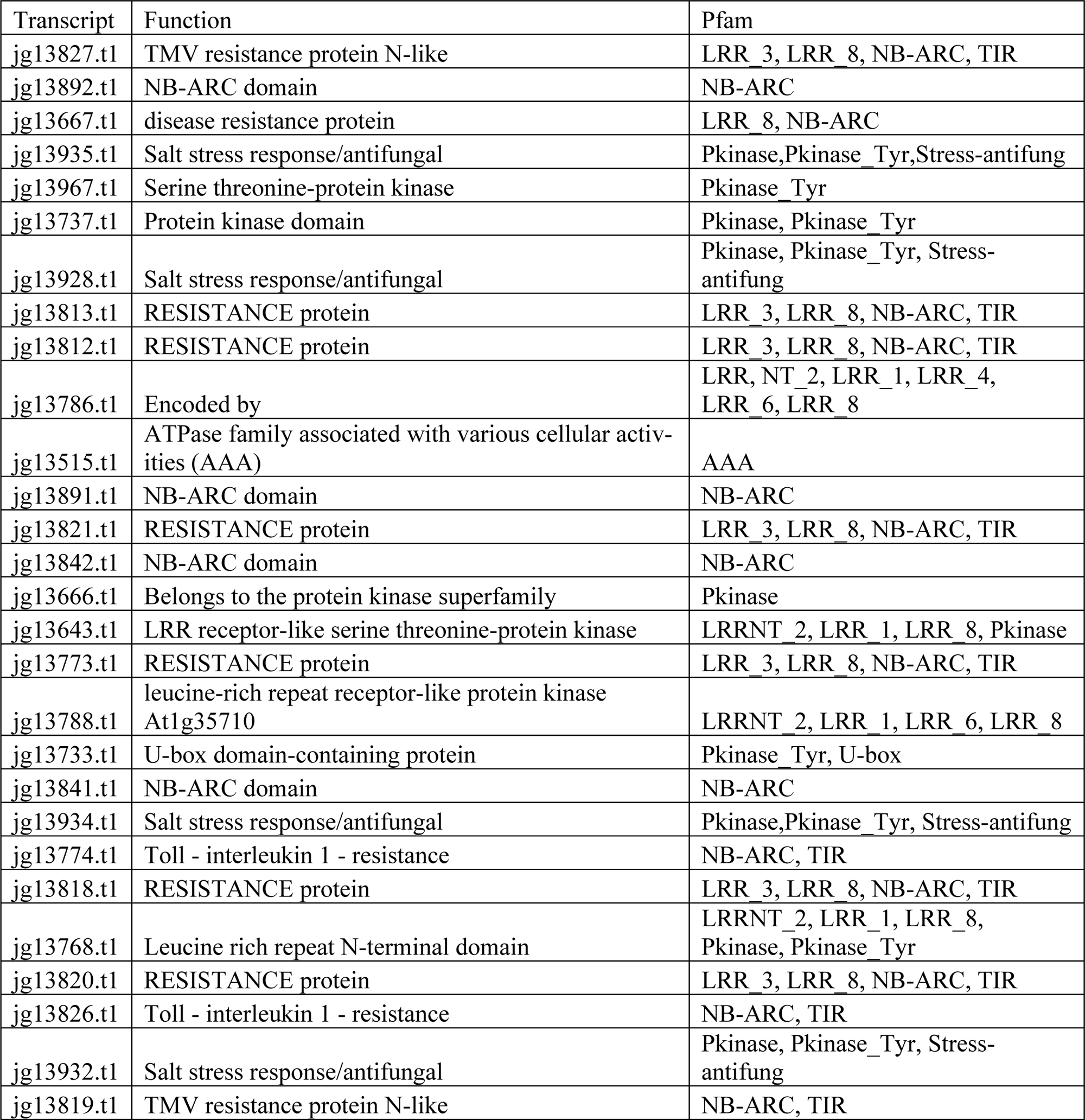
Functional annotation of *R* gene transcripts identified adjacent to the CRKN resistance markers in the SB22 genome.

## Discussion

Here, we present the chromosome-level assembly of *S. bulbocastanum* (selection 22). The present genome assembly of the *S. bulbocastanum* (SB22) genome offers a more complete chromosome-level assembly than the existing assembly. The earlier assembly reported assembly completeness of 96.6%, with 93.7% single copy and 2.9% duplicated BUSCOs compared to 95.7 % of our assembly, with 90.2 % single copy and 5.5 % duplicated BUSCOs. Relative genome completeness according to these metrics was comparable but not equal to DM6.1, which is 97.9 % (Pham et al. 2020). The LAI index of 8.73 of SB22 is lower compared to other high-quality genomes like DM 6.1 which has an LAI score of 13.56 (Pham et al. 2020). The highly accurate genomes of rice and maize have LAI scores of 21.71 and 20.7 respectively (Ou *et al*. 2018; Wu *et al*. 2023). This assembly of SB22 is comparable if not better than the existing assemblies of *Solanum* species. A comparison of the SB22 assembly with DM6.1 revealed several inversions in the genome. The incompatibility of cultivated tetraploid potatoes with *S. bulbocastanum* when crossing by traditional methods is well documented. *S bulbocastanum* exhibited visible variations during cell division and structural variations in the genome (Hermsen and Ramanna 1976; Jackson and Hanneman 1999). These structural variations agree with an earlier linkage map study indicating the presence of inversions in chromosomes of the *Solanum bulbocastanum* genome compared to *S. tuberosum* and tomato genomes (Iorizzo *et al*. 2014).

The repeat landscape in SB22 is similar to that of the DM6.1 genome, in which 66.8% (495.7Mb) consisted of repeat elements (Pham et al.2020). Retroelements and unclassified repeats represented similar proportions of repeat elements in DM6.1 (36.83 and 25.81 %) and SB22 (31.63% and 27.64 %) genomes. The slightly higher proportion of unclassified repeats in SB22 relative to DM6.1 which may be due to the quality of repeat libraries used. Tang *et al*. (2022) published a contig-level assembly of *S. bulbocastanum* which had a monoploid genome size of 666 Mb with 301 contigs and a repeat content representing 59 % of the genome. Microsatellites are evenly distributed in plant genomes; they are useful in plant breeding programs, fingerprinting and mapping studies, species discrimination, and marker-assisted selection programs. They can also be used for a variety of plant genetics applications due to their reproducibility, multiallelic nature, co-dominant inheritance, relative abundance, and good genome coverage (Remya *et al*. 2010). In the present study, we identified 62,429 microsatellites in the SB22 genome. The findings of our study are in line with the abundance of microsatellites in four other potato genomes (DM, RH, M6, and *S. commersonii*), reported earlier by Jian et al. (2021), where di-nucleotide repeats were reported to be the most abundant followed by tri-, tetra- and hepta-nucleotide repeats, with penta-nucleotide repeats being the least abundant. Our study reports ∼1 microsatellite per kb, which is higher than the earlier study, perhaps due to the parameter set used for prediction and assembly quality.

In plants, defense against parasitic organisms depends on the interaction between pathogen avirulence gene loci (*avr*) and corresponding plant disease resistance (*R*) loci. In the SB22 genome, we mapped 2,310 ‘*R’* genes, including CNL and TNL genes that may play a role in disease resistance. The SB22 genome has a lower number of *R* genes than DM6.1. The tetraploid nature, genome duplication events, and larger genome size of DM6.1 may explain the higher number of *R* genes in the DM6.1 genome. Polyploidization and genome duplication led to the expansion of *R* genes in tetraploid potatoes (Xu *et al*. 2011). We also note that the number of *R* genes varies greatly among species of potatoes, ranging from 478 in *Solanum morelliforme* to 1,976 in *Solanum chacoense*; indicating diverse evolutionary histories (Tang *et al*. 2022). The number of TNL genes identified in DM6.1 is, by our analysis, higher than the CNL genes. This finding is in line with an earlier report (Jupe *et al*. 2012), but the number of CNL genes in SB22 was greater than the number of TNL genes. This could be due to the SB22gene models being of lower quality than the well-established, high-quality gene models of DM6.1. The number of RLP and RLK genes was higher in DM6.1 (394 and 388) compared to their numbers in SB22 (271 and 304). These genes confer apoplastic immunity and provide a broader spectrum of resistance. Prodhomme et al. 2020 identified RLK and RLP genes as candidates for various pathotypes of potato wart resistance. A study on lysine motif receptor-like kinases (LysM-RLKs) in potatoes identified 77 RLKs, of which 10 had at least one lysine motif (LysM). LysM containing RLKs are known to play an important role in the perception of elicitors or pathogen-associated molecular patterns (PAMPs) in plants (Nazarian-Firouzabadi *et al*. 2019). The RLK and RLP identified in SB22 will aid in the elucidation of non-NLR resistance sources, subsequently add to breeding program efficiency. In a previous study, Bali et al. (2022) identified markers associated with CRKN resistance. We mapped those markers to chromosome 11 of the assembled SB22 genome and predicted the probable *R* genes contributing to CRKN resistance. *R* gene prediction may vary with the genome assembly quality, evidence, model, and accuracy. Hence, more advanced sequencing approaches, like PacBio HiFi, SB22 tissue-specific RNAseq reads, and more accurate *R*-gene prediction models, can be considered for future updates to this genome.

## Conclusion

Drafting the genome of SB22 *Solanum bulbocastanum* has significant implications for breeding programs aimed at improving disease resistance in related crops. This species is a source of resistance to various pests and diseases, including Columbia root-knot nematode and late blight caused by the pathogen *Phytophthora infestans*. The availability of this genomic resource will facilitate downstream discovery of novel resistance genes and accelerate the breeding of crops with improved disease resistance. *R-*genes, which may be responsible for CRKN resistance, were also found in the region of mapped CRKN resistance markers in SB22. We identified two TNL genes in close proximity to the CRKN resistance markers SB_MC1chr11-INDEL-4 and SB_MC1chr11-SSR20. By identifying and characterizing genes responsible for disease resistance in *S. bulbocastanum*, breeders can predictably and rapidly introgress these genes into elite cultivars through molecular approaches such as marker-assisted selection or genomic selection. The SB22 genome and its gene annotation set also hold potential for uncovering genes related to agronomic quality and potato processing traits, which may be of interest for future applications.

## Data Availability Statement

The authors confirm that the data supporting the findings of this study are available within the article [and/or] its supplementary materials.

## Conflict of Interest

The Authors Declare that there is no conflict of interest.

## Funder Information

We thank The Northwest Potato Research Consortium and Oregon State University startup fund for Dr Vidyasagar Sathuvalli for funding and support for this project.

---

**Table S1:** Functional annotation of *R* gene transcripts identified by Drago3 containing CNL and TNL Domains.

| Transcript | Description | PFAMs |
| --- | --- | --- |
| jg23482.t1 | Toll - interleukin 1 - resistance | LRR_8, NB-ARC, TIR |
| jg23481.t1 | Toll - interleukin 1 - resistance | LRR_8, NB-ARC, TIR |
| jg23470.t1 | Toll - interleukin 1 - resistance | LRR_8, NB-ARC, TIR |
| jg23459.t1 | TMV resistance protein N-like | NB-ARC, TIR |
| jg27897.t1 | NB-ARC domain | NB-ARC |
| jg25572.t1 | Belongs to the disease resistance NB-LRR family | NB-ARC |
| jg27833.t1 | TMV resistance protein N-like | NB-ARC, TIR |
| jg26680.t1 | disease resistance | NB-ARC |
| jg21449.t1 | Belongs to the disease resistance NB-LRR family | NB-ARC |
| jg37222.t1 | late blight resistance protein homolog | NB-ARC |
| jg36558.t1 | disease resistance protein | LRR_8, NB-ARC |
| jg34386.t1 | Belongs to the disease resistance NB-LRR family | NB-ARC, RPW8 |
| jg35694.t1 | RESISTANCE protein | LRR_3, NB-ARC |
| jg37852.t1 | resistance protein | LRR_8, NB-ARC, TIR |
| jg37851.t1 | ATPase family associated with various cellular activities (AAA) | LRR_1, LRR_8, NB-ARC, TIR |
| jg3142.t1 | NB-ARC domain | NB-ARC |
| jg33001.t1 | Resistance protein | NB-ARC |
| jg32391.t1 | Disease resistance protein | LRR_8, NB-ARC, TIR |
| jg32951.t1 | Resistance protein | NB-ARC |
| jg4977.t1 | NB-ARC domain | LRR_8, NB-ARC |
| jg4770.t1 | ADP binding | LRR_3, NB-ARC, TIR |
| jg6756.t1 | TMV resistance protein N-like | NB-ARC, TIR |
| jg6751.t1 | TMV resistance protein N-like | NB-ARC, TIR |
| jg6774.t1 | Belongs to the disease resistance NB-LRR family | NB-ARC |
| jg6785.t1 | Toll - interleukin 1 - resistance | NB-ARC, TIR |
| jg494.t1 | Belongs to the disease resistance NB-LRR family | LRR_8, NB-ARC |
| jg12945.t1 | Toll - interleukin 1 - resistance | NB-ARC, TIR |
| jg13774.t1 | Toll - interleukin 1 - resistance | NB-ARC, TIR |
| jg13826.t1 | Toll - interleukin 1 - resistance | NB-ARC, TIR |
| jg10002.t1 | resistance | - |
| jg10001.t1 | resistance | - |
| jg10001.t2 | resistance | - |
| jg9993.t1 | resistance | - |
| jg9844.t1 | disease resistance protein | LRR_8, NB-ARC |
| jg8287.t1 | NB-ARC domain | LRR_4, LRR_8, NB-ARC |

**Figure S1:**
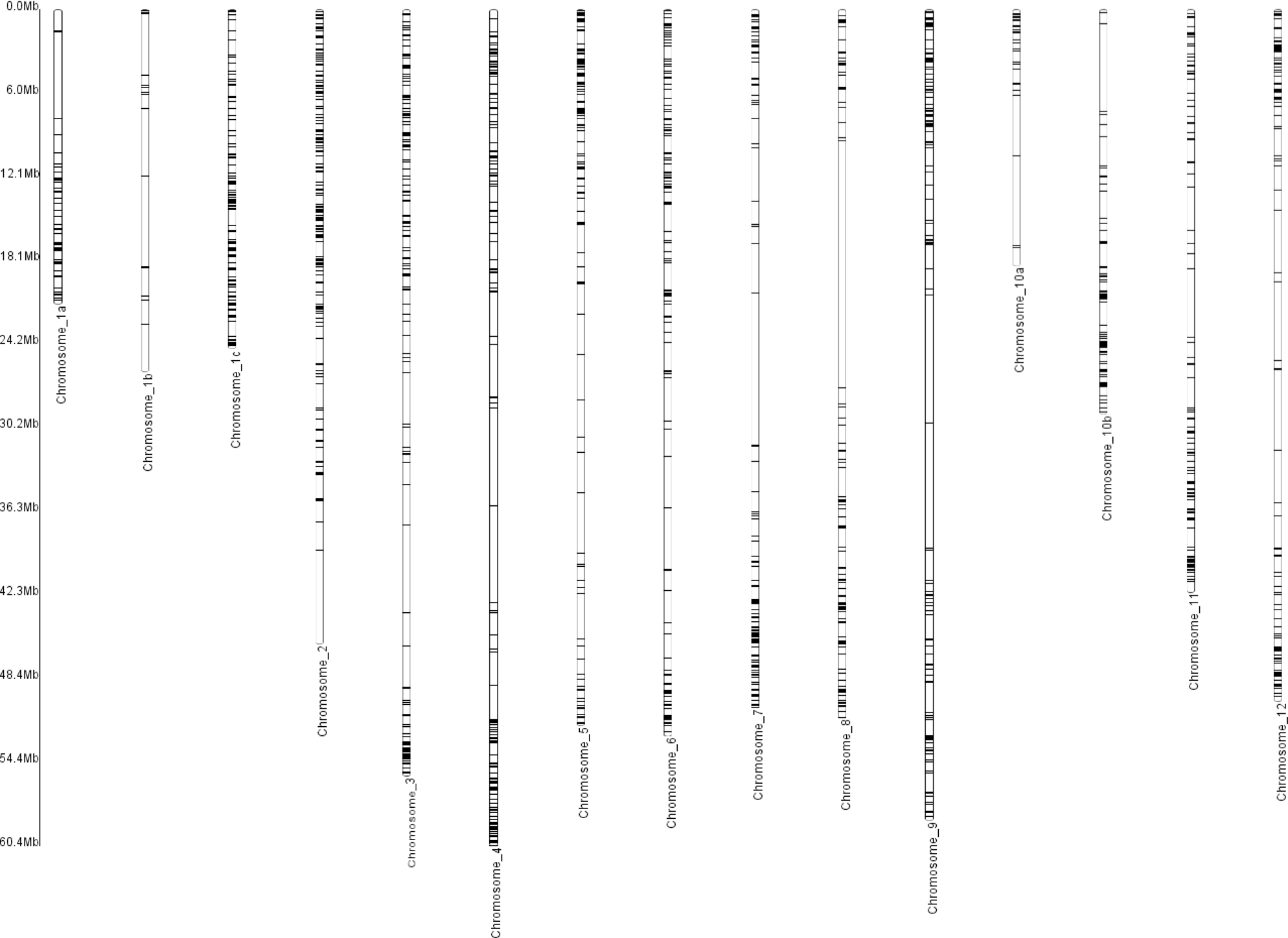
K-mer based genome size and heterozygosity estimation using *Solanum bulbocastanum* raw reads.

**Figure S2:**
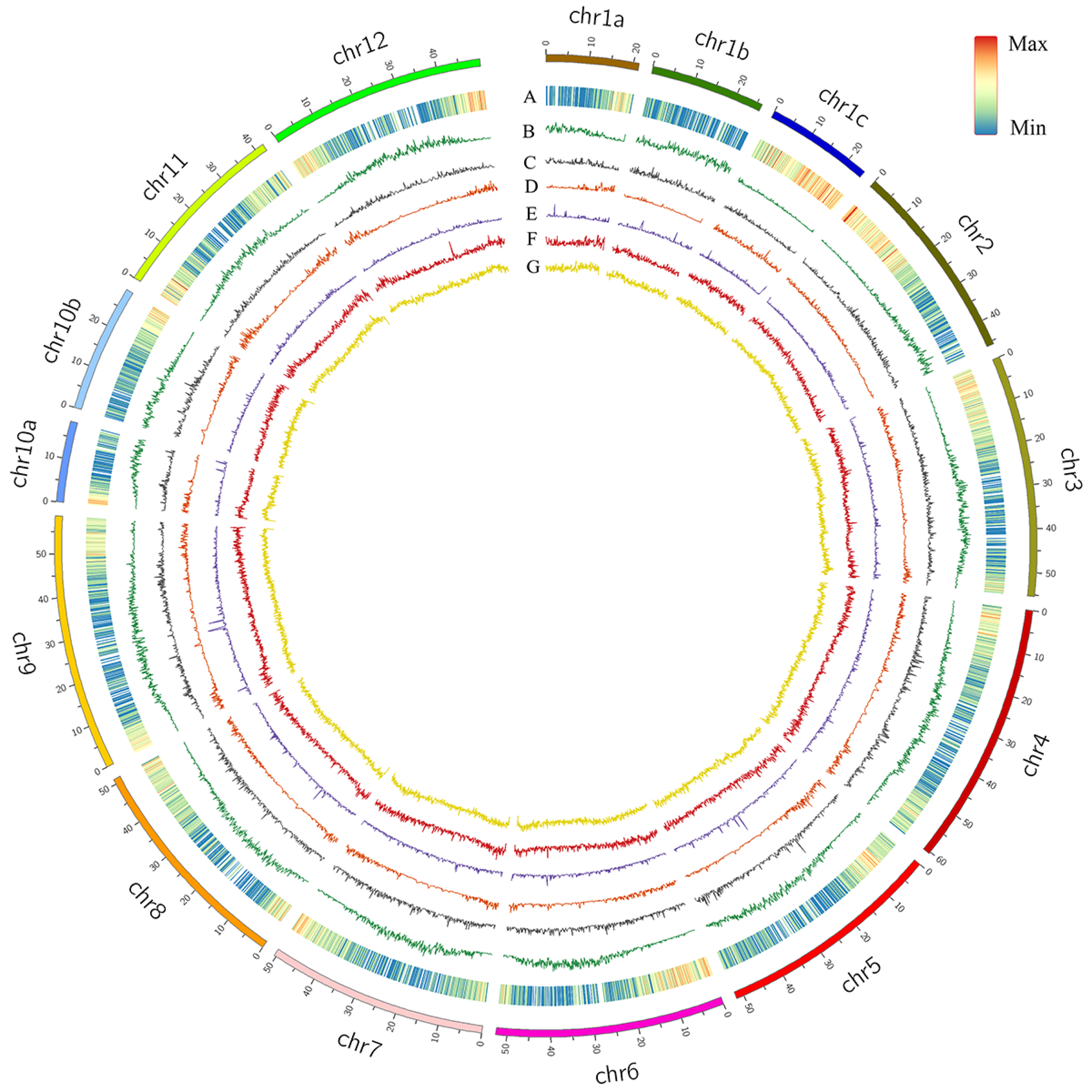
Positions of R genes identified through Drago3 across 15 scaffolds (12 chromosomes) of the SB22 genome.

